# STK19 is a transcription-coupled repair factor that participates in UVSSA ubiquitination and TFIIH loading

**DOI:** 10.1101/2024.07.17.604011

**Authors:** Yuanqing Tan, Meng Gao, Yanchao Huang, Delin Zhan, Sizhong Wu, Jiao An, Xiping Zhang, Jinchuan Hu

**Affiliations:** Shanghai Fifth People’s Hospital, Fudan University, and Shanghai Key Laboratory of Medical Epigenetics, International Co-laboratory of Medical Epigenetics and Metabolism (Ministry of Science and Technology), Institutes of Biomedical Sciences, Fudan University, Shanghai 200032, China

## Abstract

Transcription-coupled repair (TCR) is the major pathway to remove transcription-blocking lesions. Although discovered for nearly 40 years, the mechanism and critical players of mammalian TCR remain unclear. STK19 is a factor affecting cell survival and recovery of RNA synthesis in response to DNA damage, however, whether it is a necessary component for TCR is unknown. Here we demonstrated that STK19 is essential for human TCR. Mechanistically, STK19 is recruited to damage sites through direct interaction with CSA. It can also interact with RNA polymerase II *in vitro*. Once recruited, STK19 plays an important role in UVSSA ubiquitination which is needed for TCR. STK19 also promotes TCR independent of UVSSA ubiquitination by stimulating TFIIH recruitment through its direct interaction with TFIIH. In summary, our results suggest that STK19 is a key factor of human TCR that links CSA, UVSSA ubiquitination and TFIIH loading, shedding light on the molecular mechanisms of TCR.

## Introduction

Bulky adducts induced by various exogenous factors including UV, cisplatin, benzopyrene and aflatoxin as well as endogenous agents such as formaldehyde pose a formidable threat to cells, as they can interfere with DNA replication and transcription, leading to mutations and cell death(1,2). The main mechanism for mammals including human to eliminate these damages is nucleotide excision repair (NER), which can be divided into global genome repair (GGR) and transcription-coupled repair (TCR) based on distinct modes of damage recognition(1). GGR recognizes damage-caused double strands distortion through the coordinated action of DDB2 and XPC(3). In contrast, TCR is initiated by elongating RNA polymerase II (PolII) blocked by lesions. Stalled PolII recruits repair factors including CSB, CSA and UVSSA(4). While significant progress has been made in understanding human GGR, the detailed molecular mechanisms underlying TCR remain elusive due to the absence of an *in vitro* system(5). Briefly, when an elongating PolII is blocked by a lesion, CSB is recruited and its interaction with PolII is enhanced by ARK2N-CK2-mediated phosphorylation(6). Then CSB can push the stalled PolII to overcome “small” lesions with its translocase activity(7,8). If the lesion is too “big” to be bypassed, CSA, in the form of the ubiquitin E3 ligase CRL4^CSA^ (CSA-DDB1-Cul4A-Rbx1), will be recruited to ubiquitinate surrounding factors including PolII and CSB(9). Then UVSSA will be loaded in complex with the deubiquitinase USP7(10–13). These repair factors, together with stalled PolII, can recruit TFIIH to complete damage recognition(14). In both GGR and TCR, the TFIIH complex is loaded as a scaffold, followed by the recruitment of XPA and RPA as well as the endonucleases XPF-ERCC1 and XPG which excise the damaged segments on 5’ and 3’ ends, respectively(3). The resulting gaps are then sealed by DNA polymerases and ligases to restore the integrity of the double strands(3,5).

Classical TCR factors, including CSA, CSB and UVSSA, are identified through the studies of human genetic disorders such as Cockayne syndrome (CS) and UV-sensitive syndrome (UV^S^S)(4,15). However, potential additional participants in TCR unrelated to human diseases remain unclear. Recent advances in proteomics and CRISPR screen have uncovered novel players in TCR, such as the ubiquitination of PolII catalytical subunit RPB1 at the K1268 residue and the transcription elongation factor ELOF1(16–19). Although it has long been known that RPB1 is ubiquitinated upon UV irradiation, K1268 was recently identified as the major site for this modification by proteomics studies. RPB1-K1268 ubiquitination is directly involved in UV-induced PolII degradation and UVSSA ubiquitination, which is crucial for the regulation of PolII pool and TCR, respectively(18,19). Additionally, ELOF1 was recently identified through both CSB interactome and CRISPR screens. Unlike other classical TCR factors, ELOF1 always associates with elongating PolII even without damage(16,17). Loss of ELOF1 can prevent the ubiquitination of RPB1 and UVSSA to inhibit TFIIH loading and thus TCR(20). Intriguingly, no reported mutations of RPB1-K1268 or *ELOF1* are associated with human genetic diseases.

Another new TCR factor candidate, STK19, which is a putative protein kinase, has emerged from genome-wide screens and proteomics studies(21,22). Moreover, its loss compromised the recovery of RNA synthesis after UV irradiation(16). However, direct evidence for the involvement of STK19 in TCR is currently lacking. Notably, STK19 exists in two isoforms, the 364aa isoform (STK19_L_) and the 254aa isoform (STK19_S_) (Figure1A). The D89N mutation on STK19_L_ has been implicated as a gain-of-function mutation in melanoma progression depending on its kinase activity(23), although debates surround the existence of STK19_L_ *in vivo* as well as its kinase activity(24,25). The specific isoform and the role of STK19’s putative kinase activity in its DNA damage-related functions also remain unclear.

**Figure 1.**
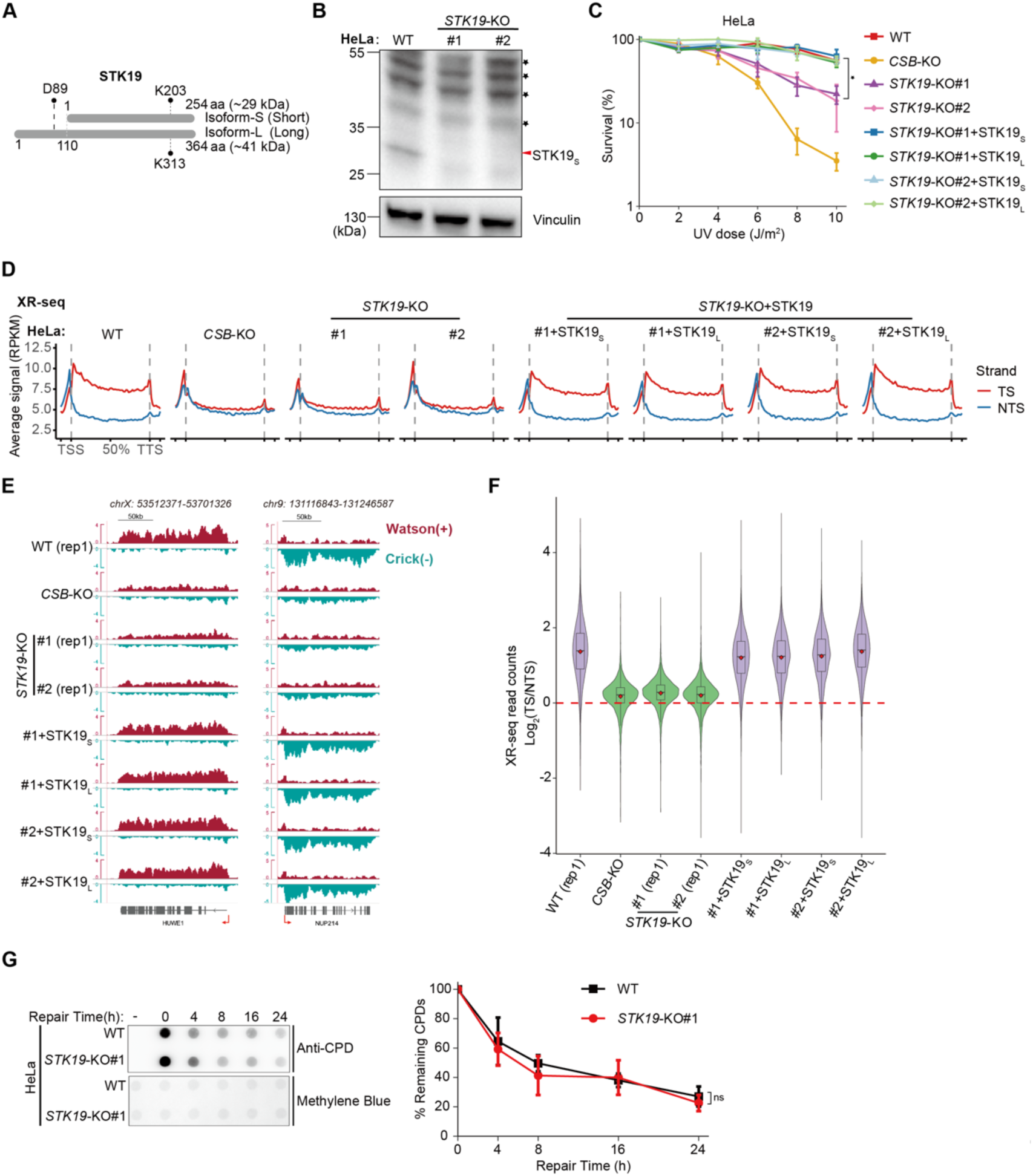
STK19 is essential for transcription-coupled repair in HeLa cells. (A) Schematic representation of two human STK19 protein isoforms. D89 in the long isoform is the mutation hotspot in melanoma, while K203 in the short isoform or K313 in the long isoform is the key residue for the putative kinase activity(23). (B) Western blot analysis showing the knockout of STK19 in HeLa cells. Non-specific bands are indicated with black asterisk. (C) UV survival of HeLa WT, *CSB-KO*, *STK19*-KO and rescued cells measured by clonogenic assay. The data are presented as the means ± SEMs (n = 3, \**P* < 0.05). (D) Metaplots of XR-seq signal from HeLa cell lines. A total of 9973 nonoverlapping protein-coding genes were used. RPKM, reads per kilobase per million mapped reads; TSS, transcription start site; TTS, transcription terminal site; TS, template strand; NTS, non-template strand. (E) IGV snapshot showing XR-seq signals in two representative genes. (F) Relative quantification of TCR activity based on the ratio of XR-seq read counts from TSs and NTSs in expressed protein-coding genes (n = 8600, TPM > 5). (G) Dot-blot assay of CPD damage for assessing global genome repair activity in HeLa WT and *STK19*-KO cells. Representative images are shown (Left). The quantification data are presented as the means ± SEMs (n = 3, ns: not significant) (Right). Results of replicate 1 for WT and KO cells are shown in (D-F). See also Supplementary Figure S1 and S2.

In this study, we demonstrate that STK19 is a *bona fide* TCR factor in human cells. Its deletion can inhibit TCR but not GGR, and complement of either isoform of STK19 can rescue TCR. However, the putative kinase activity is not required for repair. Unlike ELOF1 that is a component of transcription elongation complex, STK19 is recruited to damage sites through its interaction with CSA. Moreover, STK19 plays important roles in UVSSA mono-ubiquitination and TFIIH loading to support TCR. Research on STK19 will help complete a crucial piece of the puzzle in understanding the molecular mechanism of human transcription-coupled repair.

## MATERIALS AND METHODS

### Cell lines and cell culture

The HeLa-S3 cells were purchased from American Type Culture Collection. XP-C (XP4PA-SV-EB, GM15983) mutant human skin fibroblasts were purchased from Coriell Institute. Cells were maintained in standard Dulbecco’s modified Eagle’s medium (DMEM, Gibco) supplemented with 10% fetal bovine serum (FBS, Royacel) and 1% penicillin–streptomycin (Gibco) in a humidified incubator at 37 °C and 5% CO_2_.

### Generating mutant cell lines by CRISPR-Cas9 technology

*CSB*-KO, *CSA*-KO, *UVSSA*-KO and RPB1-K1268R mutated XP-C cell lines have been described previously(26). For generating other KO cells, HeLa or XP-C cells were transiently transfected with pX330-mCherry (Addgene, 98750) or pX459-puromycin (Addgene, 62988) plasmid(27) containing appropriate sgRNAs using HighGene transfection reagent (ABclonal). Transfected cells were FACS sorted or selected using 1μg ml^−1^ puromycin (Selleck) for 1 day, and single cells were seeded by limiting dilution in 96-well plates to allow expansion. Isolated KO clones were verified by Sanger sequencing of genomic DNA and/or western blot. The sgRNA sequences are presented in Supplementary Table S1.

### Generation of stable cell lines

For the generation of STK19 complemented or overexpressed cell lines, the wild-type human STK19_L_ (364aa isoform) and STK19_S_ (254aa isoform) cDNA were cloned into pMXs-MCS-3xHA-IRES2-Puro retroviral expression vector (a gift from Dr. Feilong Meng, Chinese Academy of Sciences, Shanghai, China) by Gibson Assembly Kit (Vazyme), respectively. The amino acid substitution mutants, kinase-dead STK19 (S-K203P/L-K313P), were generated from the wild-type STK19-encoding plasmid by site-directed PCR mutagenesis kit (Beyotime) and verified using Sanger sequencing. For virus production, HEK-293T cells were co-transfected with STK19-encoding plasmids constructed above together with packaging plasmid PCL10A1 (a gift from Dr. Feilong Meng) using HighGene transfection reagent. Viral particles were collected 48 h after transfection, filtered through 0.45-μm filters, and infected into *STK19*-KO or other cell lines. Infected cells were selected by 2 μg ml^−1^ puromycin, then verified by western blotting.

For the construction of XPD or XPB overexpressed cell lines, pMXs-MCS-3xHA-IRES2-Puro retroviral expression vector carrying the wild-type human XPD or XPB cDNA and packaging plasmid PCL10A1 were co-tranfected into HEK-293T cells to produce virus. The harvested virus was then infected into XP-C WT, *CSA*-KO and *STK19*-KO cells, respectively. Infected cells were selected by 2 μg ml^−1^ puromycin, then verified by western blotting.

For the generation of UVSSA-WT or UVSSA-K414R complemented cell lines, the wild-type human UVSSA or UVSSA-K414R cDNA were cloned into pCDH-3xFlag-MCS-IRES2-Blast-EF1-copGFP plasmid. For virus production, HEK-293T cells were co-transfected with UVSSA- or UVSSA-K414R-encoding plasmid together with packaging plasmid pMD2.G (Addgene, #12259) and psPAX2 (Addgene, #12260) using HighGene transfection reagent. Viral particles were collected 48h after transfection, filtered through 0.45-μm filters, and infected into XP-C UVSSA-KO cell line. Infected cells were selected by 5 μg ml^−1^ Blasticidin (Selleck), then verified by western blotting. Sequences of all primers are presented in Supplementary Table S2.

### RNA interference

For STK19 knockdown, siRNA transfection was performed 2 days before each experiment using Lipofectamine RNAiMAX (Thermo Fisher Scientific) with a pool of three individual siRNAs (Sangon Biotech) against STK19_S_ according to the manufacturer’s instructions. Knockdown efficiency was determined by western blotting. The siRNA target sequences used in this study are listed in Supplementary Table S1.

### Detection of UV-induced Pol II-pSer2 ubiquitylation by Dsk2-pulldown

The detailed method has been described previously(28). Briefly, to prepare the Dsk2 affinity beads, the GST-Dsk2 proteins were expressed in *E.coli* BL21(DE3) cells transfected with pGEX-GST-Dsk2 plasmid (a gift from Dr. Fenglong Meng), and bound to glutathione agarose beads (Smart-Lifesciences). Cells were mock-treated or treated by UVC irradiation (20 J/m^2^), followed by incubation for 1 h, then collected by centrifugation. For whole cell lysates preparation, cell pellets were lysed in TENT buffer (50 mM Tris-HCl pH 7.4, 2 mM EDTA, 150 mM NaCl, 1% Triton X-100) containing protease and phosphatase inhibitors (Roche), sonicated briefly and centrifuged to discard debris. Dsk2-coated beads were added to the whole cell extracts to enrich the ubiquitylated proteins. The Dsk2-coated beads were washed, then suspended with SDS protein loading buffer (Beyotime), boiled at 98 °C for 5 min and centrifuged. The supernatants were collected and analyzed by western blotting.

### UV survival assay

For HeLa cell lines, 1,000 cells were seeded in triplicated 6-well plates. In the following day, cells were treated with UVC irradiation (0-10 J/m^2^ as indicated). Following treatment, cells were grown for 7-10 days. After removing culture medium and washing with 1x PBS, cells were fixed with 100% methanol for 10 min at room temperature and stained with 0.5% (w/v) crystal violet (Sangon Biotech) solution in 25% methanol for 10 min at room temperature. After washing with water, colonies were counted using Image J software.

For XP-C cell lines, the UV survival assay was measured using Alamar blue assay(29,30). Briefly, 1,000 cells were seeded in quadruplicate in 24-well plates and treated in the next day with UVC irradiation (0,1,2,4 J/m^2^). After five days, growth medium was removed and replaced with 0.5 ml Alamar blue cell viability reagent (36 μg ml^−1^ resazurin (Sigma) in phosphate-buffered saline (PBS)), then plates were incubated for 2 h at 37 °C. Viability was assessed by using fluorescence (560 nm excitation/590 nm emission). Technical quadruplicates were averaged and treated as one biological replicate.

### Dot blot assay

Equal numbers of HeLa cells were seeded in 60mm dishes, and treated with UV irradiation (UVC, 20J/m^2^) in the next day, followed by different incubation time as indicated to allow for repair. Cells were collected and saved at -80°C until genomic DNA (gDNA) extraction. Dot blot assay was performed as previously reported(31). Briefly, gDNA was extracted using a DNA extraction kit (Thermo Fisher Scientific). DNA was quantified with Qubit dsDNA HS Assay Kit (Thermo Fisher Scientific) and diluted to 1 ng μL^-1^. Diluted DNA was boiled for 10min at 100 °C and immediately placed on ice. Three Whatman paper (Cytiva) presoaked in 6x SSC (20x SSC: 175.3g NaCl and 88.2g sodium citrate in 1L sterile water with pH 7.0) for 10 min, and the positively charged nylon membrane (Cytiva) presoaked in sterile water for 5min and 6x SSC for 10 min was placed in a BIO-RAD dot blot apparatus. Membrane was washed once with 400μL/well TE buffer (10mM Tris, 1mM EDTA, pH 8.0). Denatured DNA (300μL) was loaded on the nylon membrane, and 400μL/well 2x SSC was used to wash the membrane. DNA was fixed in a vacuum drier at 80°C for 2h. The membrane was washed in TBST buffer (TBS with 0.05% Tween 20) for 10min, and blocked with blocking solution (5% non-fat powered milk in TBST) for 1h. After washing 3 times with TBST, the membrane was incubated overnight at 4°C with anti-CPD antibody (Cosmo Bio) 1:5000 diluted in TBST with 5% BSA and 0.02% NaN_3_. Membrane was washed 4 times with TBST followed by incubation with HRP-labled anti-mouse secondary antibody (Beyotime) 1:5000 diluted in blocking buffer for 1h at room temperature. After washing another 4 times with TBST, CPDs were detected using enhance chemiluminescence reagent (Tanon). As a loading control, the membrane was stained with methylene blue solution (0.5 M sodium acetate, 0.1% methylene blue) for 10 min and de-stained in water for several minutes.

### Preparation of total chromatin fraction

One million cells were seeded for each condition (mock or UV treatment) in a 6-cm dish. In the following day, cells were mock-treated or irradiated with UV-C (40 J/m^2^), followed by 0.5 h incubation. The collected cell pellets were resuspended in lysis buffer 1 (50 mM HEPES-KOH pH 7.4, 1 mM EDTA pH 8.0, 140 mM NaCl, 0.25% Triton X-100, 0.5% CA-630, 10% glycerol, phosphatase and protease inhibitor cocktail). After incubation on ice for 20 min, the lysed cells were centrifuged for 3 min at 20,000g at 4 °C, followed by removal of the supernatants. The harvested chromatin pellets were sequentially washed with lysis buffer 1 and lysis buffer 2 (10 mM Tris-HCl pH 8.0, 1 mM EDTA pH 8.0, 200 mM NaCl and 0.5 mM EGTA pH 8.0), followed by centrifugation and removal of the supernatants. The chromatin pellets were then fragmented with lysis buffer 1-IP (50 mM HEPES-KOH pH 7.4, 140 mM NaCl, 0.25% Triton X-100, 0.5% CA-630, 10% glycerol, phosphatase and protease inhibitor cocktail) supplemented with 500 U/ml Super Nuclease (Beyotime) for 15 min on ice. The fragmented chromatin was dissolved by adding the SDS protein loading buffer (Beyotime) and boiling for 10 min at 100 °C. The samples were analyzed by western blotting.

### Western blot

For whole cell extracts, cell pellets were resuspended in RIPA buffer (10 mM Tris-HCl pH 8.0, 1 mM EDTA pH 8.0, 140 mM NaCl, 1% Triton X-100, 0.1% SDS and 0.1% Na-DOC) on ice for 10 min, and briefly sonicated using a Q800 Sonicator (Qsonica). The samples were then centrifuged at 20,000g for 5 min at 4 °C to discard debris. Collected lysates were denatured by addition of SDS protein loading buffer and boiling at 100 °C for 10 min. Samples were resolved by 4-12% or 4-20% gradient-MOPS-SDS-PAGE (ACE Biotechnology) or gradient-Tris-Gly-SDS-PAGE gels (Beyotime). Proteins were transferred to nitrocellulose membranes (PALL), followed by blocking for 1 h at room temperature in 5% skim milk in TBST (50 mM Tris-HCl pH 7.6, 150 mM NaCl, 0.1% Tween 20). The membrane was washed 3 times in TBST and incubated with indicated primary antibodies in 5% BSA in TBST overnight at 4 °C. The membrane was washed three times in TBST, followed by incubation for 1h at room temperature with 1:5000 diluted HRP-conjugated secondary antibodies in 5% skim milk in TBST. After extensive washing with TBST, the proteins were detected using enhance chemiluminescence reagent. All primary and secondary antibodies information were listed in the Supplementary Table S3.

### XR-seq assay

XR-seq was performed based on previous ATL-XR-seq protocol(32) with minor modification. In short, HeLa cells were cultured to 80% confluency in two 15-cm plates per sample. Cells were treated with UVC (20 J/m^2^), followed by 1h incubation. The harvested cells were lysed in TENT buffer (50 mM Tris-HCl pH7.4, 2 mM EDTA, 150 mM NaCl, 1% Triton X-100) on ice for 20 min, followed by centrifugation at 20,000g at 4 °C for 20 min and collection of supernatants. The collected supernatants were treated sequentially with RNase A (Sigma) and proteinase K (NEB), followed by phenol–chloroform extraction and ethanol precipitation. Resuspended DNA samples were further purified using Zymo ssDNA purification kit (D7011) to attain short single-stranded DNA (mainly primary excision products). The samples were tailed with poly(dA) using terminal deoxynucleotidyl transferase (NEB) and dATP, followed by ethanol precipitation. Purified samples were ligated with adaptor Ad2-ATL using Instant Sticky-end Ligase Master Mix (NEB), followed by phenol–chloroform extraction and ethanol precipitation. The DNA samples containing CPD damage were enriched by CPD Damage IP and repaired with CPD photolyase as previously described(33). DNA samples were then extended with primer 30T-O3P and NEBNext Ultra II Q5 Master Mix (NEB), followed by ExoI (NEB) treatment. The following PCR amplification and purification were performed as described previously(34). Libraries were sequenced in PE150 format on an Illumina NovaSeq platform by Mingma Technologies Company. Sequences of all oligonucleotides used in library construction are presented in Supplementary Table S4.

### Damage-seq assay

For CPD Damage-seq, the detailed method has been described previously(35) with minor modification. Briefly, cells were irradiated with UVC (20 J/m^2^), followed by incubation for 0h or 8h, and harvested by centrifugation. Genomic DNA was extracted using PureLink Genomic DNA Mini Kit (Thermo Fisher Scientific), and sonicated by a Q800 Sonicator to attain DNA fragments averagely 300-600 bp in length. DNA fragments (1 μg) were used for Damage-seq library construction. DNA fragments were subject to end repair and dA-tailing with NEBNext Ultra II DNA Library Prep Kit for Illumina (NEB), ligated to Ad1 at both ends and denatured to be incubated with CPD antibody-coated beads. The beads were washed and eluted to collect the ssDNA containing the CPD damage. Purified DNA samples were extended with primer O3P and NEBNext Ultra II Q5 Master Mix, followed by ExoI treatment. The purified DNA was then denatured and ligated to adaptor Ad2 by Instant Sticky-end Ligase Master Mix. The following PCR amplification and purification were performed as described previously(35). Libraries were sequenced in PE150 format on an Illumina NovaSeq platform by Mingma Technologies Company. For Cisplatin Damage-seq, cells were treated with 200 μM cisplatin (Sigma) for 1.5 h or 8 h, harvested by centrifugation, and subjected to Damage-seq library construction as described above except for cisplatin-damage IP which was described previously(36). Sequences of all oligonucleotides used in library construction are presented in Supplementary Table S4.

### PADD-seq assay

PADD-seq was performed as described previously(37) with minor modification. Briefly, cells were cultured to 80∼90% confluency in five 15-cm plates per sample and treated with UVC (20 J/m^2^), followed by 0.5 h incubation. Cells were cross-linked with a final concentration of 1% formaldehyde (Thermo Fisher Scientific) for 10 min at room temperature followed by neutralization with a final concentration of 150 mM glycine for 5 min. Harvested cell pellets were resuspended in 6 cell pellet volumes of cold lysis buffer 1 ( 50 mM HEPES-KOH pH 7.4, 1 mM EDTA pH 8.0, 140 mM NaCl, 0.25% Triton X-100, 0.5% CA-630, 10% glycerol) supplemented with protease inhibitor (Roche) to incubate on ice for 10 min, and then centrifuged at 850g for 10 min at 4 °C to discard the supernatants. Pellets were resuspended in 6 cell pellet volumes of cold lysis buffer 2 (10 mM Tris-HCl pH 8.0, 1 mM EDTA pH 8.0, 200 mM NaCl and 0.5 mM EGTA pH 8.0) supplemented with protease inhibitor (Roche) to incubate on ice for 10 min, and centrifuged at 850g for 10 min 4 °C to collect pellets. Pellets were resuspended in 1.5 cell pellet volumes of cold 1% SDS RIPA buffer (10 mM Tris-HCl pH 8.0, 1 mM EDTA pH 8.0, 140 mM NaCl, 1% Triton X-100, 1% SDS and 0.1% Na-DOC) supplemented with protease inhibitor and SDS to a final concentration of 1.5%. The suspension was sonicated using a Q800 Sonicator at 4 °C to attain chromatin fragments averagely 300-600 bp in length. The sonicated chromatin samples were centrifuged at 20,000g for 10 min at 4 °C to collect the supernatants. Prior to immunoprecipitation, the SDS concentration of samples were diluted to 0.1% by addition of RIPA buffer without SDS. For PADD-seq of HA-tagged STK19, XPD and XPB, the samples supplemented with BSA (Sigma), tRNA (Sigma) and protease inhibitor were incubated with HA-tag antibody coupled agarose beads (Smart-Lifesciences) pre-blocked with BSA and tRNA. After incubation overnight, beads were sequentially washed with RIPA buffer, RIPA-500 buffer (10 mM Tris-HCl pH 8.0, 1 mM EDTA, 500 mM NaCl, 1% Triton X-100, 0.1% SDS and 0.1% Na-DOC), LiCl Wash buffer (10 mM Tris-HCl pH 8.0, 1 mM EDTA, 250 mM LiCl, 0.5% CA-630 and 0.5% Na-DOC) two times for each buffer, and once with 1x TE (10 mM Tris-Cl pH 8.0 and 1 mM EDTA), then eluted with direct elution buffer (10 mM Tris-HCl pH 8.8, 5 mM EDTA, 300 mM NaCl and 1% SDS) in a heating shaker at 1,500 rpm for 20 min at 65 °C. The eluted samples were treated with RNase A at 37 °C for 30 min followed by proteinase K at 55 °C for 2 h, then incubated at 65 °C overnight to reverse cross-linking. DNA samples were purified by phenol-chloroform extraction and ethanol precipitation, and the concentration was determined by Qubit dsDNA HS Assay Kits. Purified DNA samples were subjected to CPD Damage-seq as described above. PolII-CPD PADD-seq in HeLa *STK19*-KO cells with NVP-2 treatment was performed as described previously(38).

### Protein expression and purification

Human STK19_S_ coding sequence were cloned into pET-N-His-TEV-MCS vector (Beyotime) using Gibson Assembly Kit to enable expression as an N-terminal His-TEV fusion protein. Human CSA coding sequence were cloned into pET-N-GST-PreScission-MCS vector (Beyotime) using Gibson Assembly Kit to enable expression as an N-terminal GST-PreScission fusion protein. After transformation of Rosetta2 (DE3) competent *E.coli* cells, protein expression was induced by 1 mM IPTG, followed by incubation at 18 °C overnight. Harvested cell pellets were suspended in 1x TBS (Sangon Biotech) supplemented with TieChui *E.coli* Lysis Buffer (ACE biotechnology) for 10 min on ice, followed by centrifugation and collection of supernatants. For His-tagged protein, the collected supernatants were incubated with Ni-NTA agarose (Smart-Lifesciences) at 4 °C for 1 h. Agarose were washed, then eluted with 1x TBS supplemented with 500 mM imidazole. Eluted protein was pooled, concentrated and dialyzed, then checked by SDS-PAGE. For GST-tagged protein, the collected supernatants were incubated with glutathione agarose beads at 4 °C overnight. After washing with 1x TBS, GST-tag was removed by adding GST-tagged PreScission protease (Beyotime) to resins suspended in 1x TBS, followed by overnight incubation at 4 °C. Eluted untagged CSA was pooled, concentrated and dialyzed, then checked by SDS–PAGE.

### In vitro pulldown assay

To assess protein-protein interactions, 450 nM His-STK19 was incubated with 100 nM CSA, 40 nM TFIIH complex (a gift from Dr. Yanhui Xu, Fudan University, Shanghai, China) or 40 nM RNAPII complex (a gift from Dr. Yanhui Xu)(39) in TBST buffer supplemented with 1mg/ml BSA and protease inhibitor cocktail (Roche) on a rotator at 4 °C overnight. Anti-His-tag mAb (MBL) was added to the mixture, followed by incubation for 1 h at RT. Next, pre-washed protein A/G magnetic beads (Thermo Fisher Scientific) were added to half of the samples to pull down His-STK19 and co-immunoprecipitated proteins. The other half of the samples were saved as input. Beads were incubated for 1 h at RT with rotation, followed by washing 6 times with TBST buffer. Beads and input were boiled in protein loading buffer for 10min, then subjected to 4-12 % SDS-PAGE and western blot.

### Analysis of NGS data

For XR-seq data analysis, read 1 obtained in paired-end sequencing data was trimmed with cutadapt v4.4(40) to remove poly(dA) tails, potential 3’ end adaptor sequences and low sequencing quality reads. The trimmed reads were aligned to the human reference genome (hg38) with BWA v0.7.17-r1188(41) to generate SAM (Sequence Alignment/Map format) file. The SAM file containing aligned reads was then transformed into BAM format file with SAMtools v1.9.(42) Sambamba v1.0.0(43) was used to remove duplicate reads. BAM file was then converted to BED format file by BEDTools v2.30.0.(44) Reads longer than 32-nt were discarded with Linux commands for downstream analysis. The calculation and plotting of reads length distribution and dinucleotide frequencies at each position of 26-nt reads were performed with the combination of Linux commands, BEDTools v2.30.0 and custom Python scripts. For further meta-analysis, reads coverage on genome forward and reverse strand were calculated separately with BEDTools v2.30.0 and normalized with RPKM (reads per kilobase per million mapped reads) to generate forward and reverse strand BigWig files by the combination of Linux commands and bedGraphToBigWig v2.9 from UCSC. For XR-seq profiles relative to the annotated TSSs and TTSs, protein-coding genes that do not have overlapping or neighboring genes for at least 2000 bp upstream or downstream on either strand were extracted from hg38.gtf file by Linux commands and BEDTools v2.30.0. To evaluate XR-seq profiles in high expression genes in HeLa cells, BED files harboring high expression genes (TPM>5) were extracted based on RNA-seq data (ENCODE DCC accession ENCSR000CPR, ENCSR000CPQ, ENCSR000CPP, ENCSR000CQT, ENCSR000CQI, ENCSR000CQJ) and prepared non-overlapping protein-coding genes by Linux commands and BEDTools v2.9. The calculation and plotting of profiles were conducted by custom Python and R scripts. For IGV(45) screen shot, BED files containing excision product reads were converted to BAM files using BEDTools v2.9. Each BAM file was then converted to two BigWig files using function bamCoverage in deepTools2 v3.5.1(46), one only containing reads on forward strand with parameter: --normalizeUsing RPKM, --samFlagExclude 16, -- binSize 300, --smoothLength 3000, another only containing reads on reverse strand with parameter: --normalizeUsing RPKM, --samFlagInclude 16, --scaleFactor -1, -- binSize 300, --smoothLength 3000. The calculation of read counts in gene TS or NTS strand was performed using featureCounts v2.0.6(47) with BAM files containing specific reads.

For Damage-seq and PADD-seq data analysis, paired-end reads harbored Ad1 sequences at 5’ end were discarded in pairs via cutadapt v4.4. Reads were further trimmed using fastp v0.12.4(48) and then aligned to the hg38 human genome by BWA v0.7.17-r1188. The generated SAM files containing aligned reads were converted to BAM files with SAMtools v1.9. Sambamba v1.0.0 was then used to remove duplicate reads. Read 1 with mapping quality larger than 25 in remained reads was extracted using SAMtools v1.9. BAM files were then converted to BED format files by BEDTools v2.9. The damage sites and sequences were fetched using BEDTools v2.9. For CPD Damage-seq, reads in sequences containing dipyrimidines (TT, TC, CT, CC) at damage sites were held for further analysis. For cisplatin Damage-seq, reads in sequences containing d(GpG) at damage sites were held for further analysis. For downstream meta-analysis, reads coverage were calculated and treated as described above. For PADD-seq profiles relative to the annotated TSS and TTS in XP-C cell lines, highly expressed (TPM>1), protein-coding and non-overlapping genes were selected based on RNA-seq data (ENCODE DCC accession ENCSR00CUH) and hg38.gtf by Linux commands and BEDTools v2.9. The calculation and plotting of profiles were conducted by custom Python and R scripts. The calculation of read counts in genes TS or NTS strand was performed using featureCounts with BAM files containing specific reads.

### Prediction of protein-protein interaction by AlphaFold2-Multimer, DMFold-Multimer and AlphaFold3

All canonical protein sequences used in this study, including STK19_S_ (isoform 254aa), CSA, CSB, UVSSA, ELOF1 and all subunits of TFIIH, were downloaded from UniProt database. Prediction of protein-protein interaction was performed using AlphaFold2-Multimer(49) function in AlphaFold2(50) v2.3.2 deployed on a standalone server, or two online tools DMFold-Multimer(51) (https://zhanggroup.org/DMFold/) and AlphaFold3(52). The structural alignment, interface residues calculation and visualization of protein complex were performed using Pymol v2.5.7.

## Results

### STK19 is essential for efficient transcription-coupled repair

STK19 was identified as a factor influencing cellular sensitivity to UV and other transcription-blocking lesions (TBLs) in genome-wide screens(21,22). However, whether it directly participates in TCR is unknown. To answer this question, we knocked out *STK19* in HeLa cells. Since STK19 has two isoforms (Figure1A), HeLa-*STK19*-KO cell lines was generated by CRISPR-Cas9 with sgRNAs targeting common regions of both isoforms (FigureS1A). Although the antibody cannot detect endogenous STK19_L_, the disappearance of STK19_S_ was verified by Western-blot (WB) (Figure1B). As previously reported(22), loss of STK19 sensitized cells to UV irradiation, albeit to a lesser extent than *CSB*-KO cells (Figure1C, FigureS1B,C). Moreover, complement of either STK19_L_ or STK19_S_ rescued the resistance to UV (Figure1C, FigureS1D). To assess the impact of STK19 on TCR, we performed XR-seq which can measure genome-wide distribution of NER by capturing and sequencing excised fragments to determine the repair pattern of UV-induced cyclobutane pyrimidine dimers (CPDs)(32,33) (FigureS1E). Since TCR only removes lesions from template strands (TSs), XR-seq should detect more repair signals on TSs than those on non-template strands (NTSs) in TCR-proficient cells, as shown in HeLa-WT cells (Figure1D-F). In contrast, this strand asymmetry diminished in TCR-deficient *CSB*-KO and *ELOF1*-KO cells (Figure1D-F, FigureS2A,B,F,G). Strikingly, this preferable repair of TSs was also greatly reduced in both *STK19*-KO cell lines, to a similar level as the *CSB*-KO cell line (Figure1D-F, FigureS2E-G). Complement of either isoform of STK19 in both KO cell lines can recover TCR (Figure1D-F), confirming the key role of STK19 in this pathway. Intriguingly, complement of kinase-dead STK19 (S-K203P/L-K313P) also rescued TCR (FigureS2C-G), suggesting that the potential kinase activity of STK19 is not involved in TCR. In the following studies of this paper, we will focus on the short isoform of STK19.

Although XR-seq captures both GGR and TCR, it cannot quantify total repair rate. To answer whether STK19 is also involved in GGR or the common steps of NER, total repair rates of CPDs were determined by dot-blot assay. There was no significant difference between WT and *STK19*-KO cells (Figure1G), indicating that STK19 is not involved in GGR or the common steps of NER. Taken together, these results suggest that STK19 is a *bona fide* TCR factor.

To further confirm the key role of STK19 in human TCR, we knocked out *STK19* in an XPC-deficient cell line (XP4PA-SV-EB, henceforth refer to as XP-C cells) that lacks GGR (Figure2A, FigureS3A)(53). Consistent with HeLa cells, loss of STK19 increased UV sensitivity of XP-C cells, while complement of STK19_S_ could partially rescue UV resistance (Figure2B, FigureS3B). Due to the lack of GGR which results in much less excision products than HeLa cells(54), XR-seq is not a good choice for detecting TCR in XP-C cells. Therefore, we performed Damage-seq which can measure genome-wide distribution of lesions at base resolution in a strand-specific manner in XP-C cells (FigureS3C)(35,55). Efficient TCR can selectively remove damage on TSs, resulting in less damage on TSs than NTSs after a period of repair. As shown in Figure2C,D, parental XP-C cells had pronounced reduced CPD level in TSs but not NTSs, which is a typical feature of TCR in GGR-deficient cells(26). Remarkably, this strand asymmetry disappeared in both *CSB*- and *STK19*-KO cell lines, while complement of STK19_S_ in *STK19*-KO cells could partially rescue TCR (Figure2C,D). Moreover, loss of STK19 also impeded TS-specific repair of cisplatin-adducts which could be complemented by STK19_S_ (Figure2E), confirming that STK19 is a common TCR factor.

**Figure 2.**
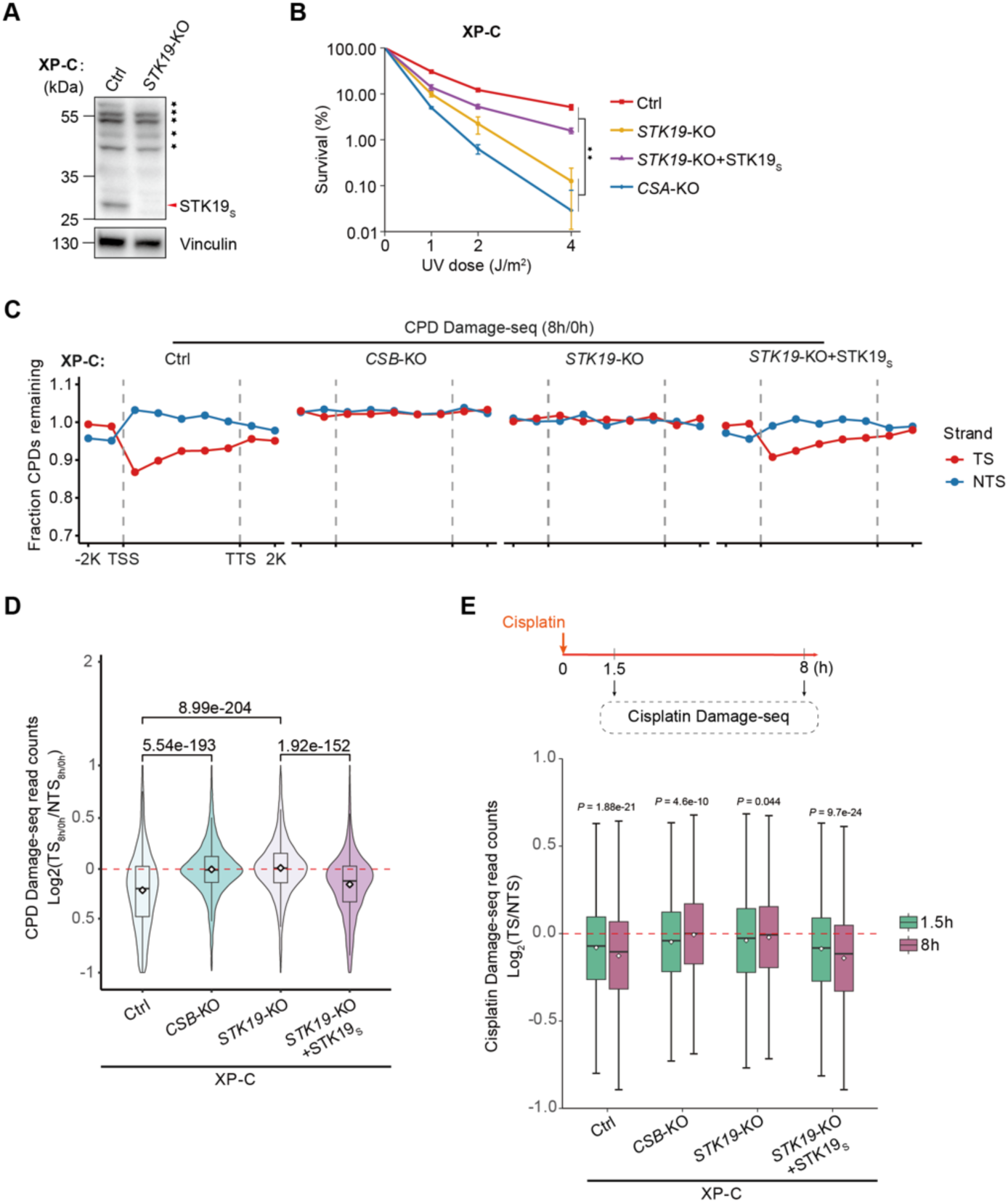
The key role of STK19 for TCR in GGR-deficient human cells. (A) Western blot analysis showing the knockout of STK19 in XP-C cells. Non-specific bands are indicated with black asterisk. (B) UV survival of parental XP-C (Ctrl), *CSA*-KO, *STK19*-KO and rescued cells measured by Alamar blue cell viability assay. The data are presented as the means ± SEMs (n = 3, \*\**P* < 0.01). (C) CPD repair measured by Damage-seq. The average fraction of CPDs remaining after 8h of repair relative to 0h was plotted along the TSs and NTSs strand for 9973 nonoverlapping protein-coding genes. (D) Relative quantification of TCR activity based on the ratio of remaining CPD in TSs to NTSs for 6946 expressed protein-coding genes (TPM > 1). (E) TCR of cisplatin damage in XP-C cells. Relative ratio of damage in TSs to NTSs based on cisplatin Damage-seq at 1.5h or 8h after cisplatin treatment in 6946 expressed protein-coding genes (TPM > 1) are shown. Fraction of remaining damage cannot be calculated since cisplatin damage is concurrently formed and repaired. TCR-proficient Ctrl and STK19-complement cells have less damage in TSs than NTSs at 1.5h, and this strand asymmetry enlarges at 8h, demonstrating the preferable repair on TSs. By contrast, the strand asymmetry of damage is less pronounced at 1.5h in both KO cells, and further reduces at 8h, indicating the lack of TCR in these cells. See also Supplementary Figure S3.

### STK19 is recruited to damage sites in a CSA-dependent manner

Classical TCR factors including CSB, CSA and UVSSA are recruited to damage sites by lesion-blocked PolII, while ELOF1, as a component of PolII elongating complex, is moving along with PolII until encountering damage(16,17). To answer whether STK19 is recruited by stalled PolII or moving along with elongating PolII even without damage, the binding of STK19 on UV-induced CPDs were measured by recently developed PADD-seq (Protein-Associated DNA Damage-sequencing) method (Figure3A)(37). It detects the distribution of DNA lesions on protein-bound DNA fragments by the combination of Chromatin immunoprecipitation (ChIP) and Damage-seq, thus can assess the direct interaction between protein and DNA damage across the genome. For TCR factors, since TCR can only deal with damage on TSs, PADD-seq should detect much higher signals on TSs than NTSs if they are recruited to damage sites. As ChIP-grade STK19 antibody was not available, ectopic expressed HA-tagged STK19_S_ and anti-HA antibody were used for PADD-seq. In the XP-C STK19-complement cells, STK19-CPD PADD-seq signal was enriched on TSs at 0.5 h after UV irradiation (Figure3B), indicating efficient recruitment of STK19 to CPDs on TSs. HA-tagged STK19_S_ was expressed in parental and TCR-deficient XP-C cells to assess the recruitment of STK19 (FigureS4). The preferred binding of STK19 on TSs nearly disappeared in *CSB*-KO and *CSA*-KO cells (Figure3C,D), suggesting that STK19 was not recruited to CPDs in these cells. In contrast, loss of ELOF1, UVSSA or XPA, as well as cells bearing RPB1-K1268R mutation that could not be ubiquitinated after UV treatment(18,19), did not abrogate the STK19-CPD interaction on TSs (Figure3C,D). These results imply that STK19 is recruited to damage sites following the recruitment of CSA during TCR rather than moving along with elongating PolII regardless of damage.

**Figure 3.**
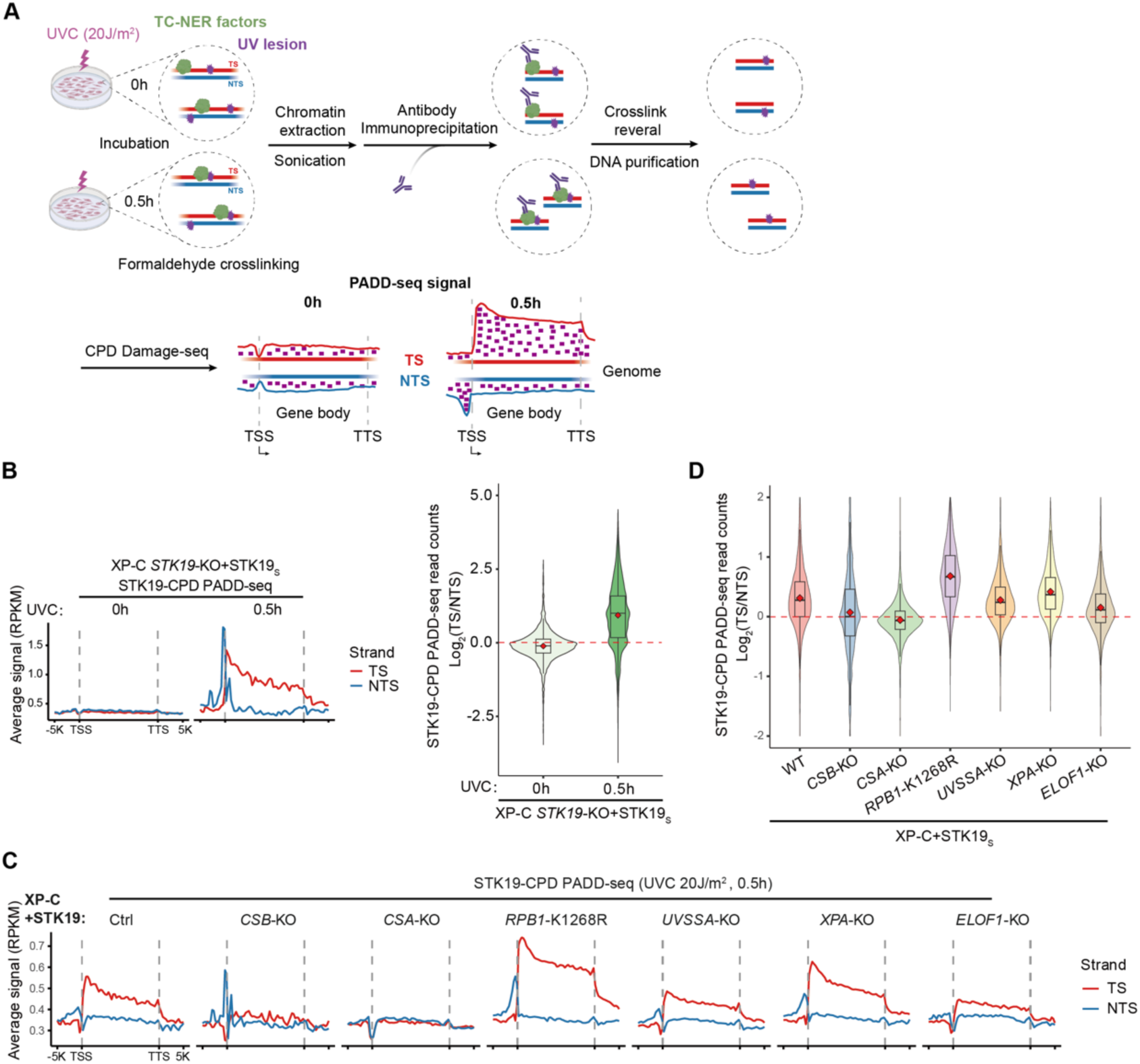
STK19 is recruited to damage sites in a CSA-dependent manner. (A) Schematic of experimental design of PADD-seq. The small peak on NTS upstream of TSS is due to divergent transcription on NTSs. (B) Validation of PADD-seq with ectopic expressed HA-tag protein. Meta-analysis of STK19-CPD PADD-seq in XP-C *STK19*-KO rescued cells at indicated time points after UVC irradiation (20 J/m^2^) is shown. A total of 4869 expressed nonoverlapping protein-coding genes were used for metaplots (left). A total of 4858 expressed protein-coding genes were used for quantification (right). (C) Metaplots of STK19-CPD PADD-seq signals in indicated XP-C cell lines overexpressing STK19_S_. A total of 4869 expressed nonoverlapping protein-coding genes were used. (D) Relative quantification of STK19-CPD PADD-seq. A total of 5381 expressed protein-coding genes were used. The number of genes is different from (B) since genes with 0 read on either strand should be discarded. See also Supplementary Figure S4.

Therefore, it is reasonable to speculate that STK19 is recruited by CSA. To test this hypothesis, AlphFold2/3(49,52) and DMFold(51) were used to predict protein-protein interactions between STK19 and other TCR factors. As shown in Figure4A,B and FigureS5A,B, STK19 was predicted to interact with CSA with high confidence, but not with CSB, ELOF1 or UVSSA. Consistent with this prediction, STK19 could pull down CSA *in vitro* (Figure4C, FigureS5C), implying that STK19 is recruited to damage sites via its direct interaction with CSA during TCR. Intriguingly, STK19 was predicted to interact with RPB1 as well (Figure4D, FigureS5D), which was confirmed by the fact that STK19 could pull down purified PolII complex *in vitro* (Figure4E, FigureS5E). However, STK19 was not recruited by lesion-stalled PolII in the absence of CSB or CSA *in vivo* (Figure3C,D).

**Figure 4.**
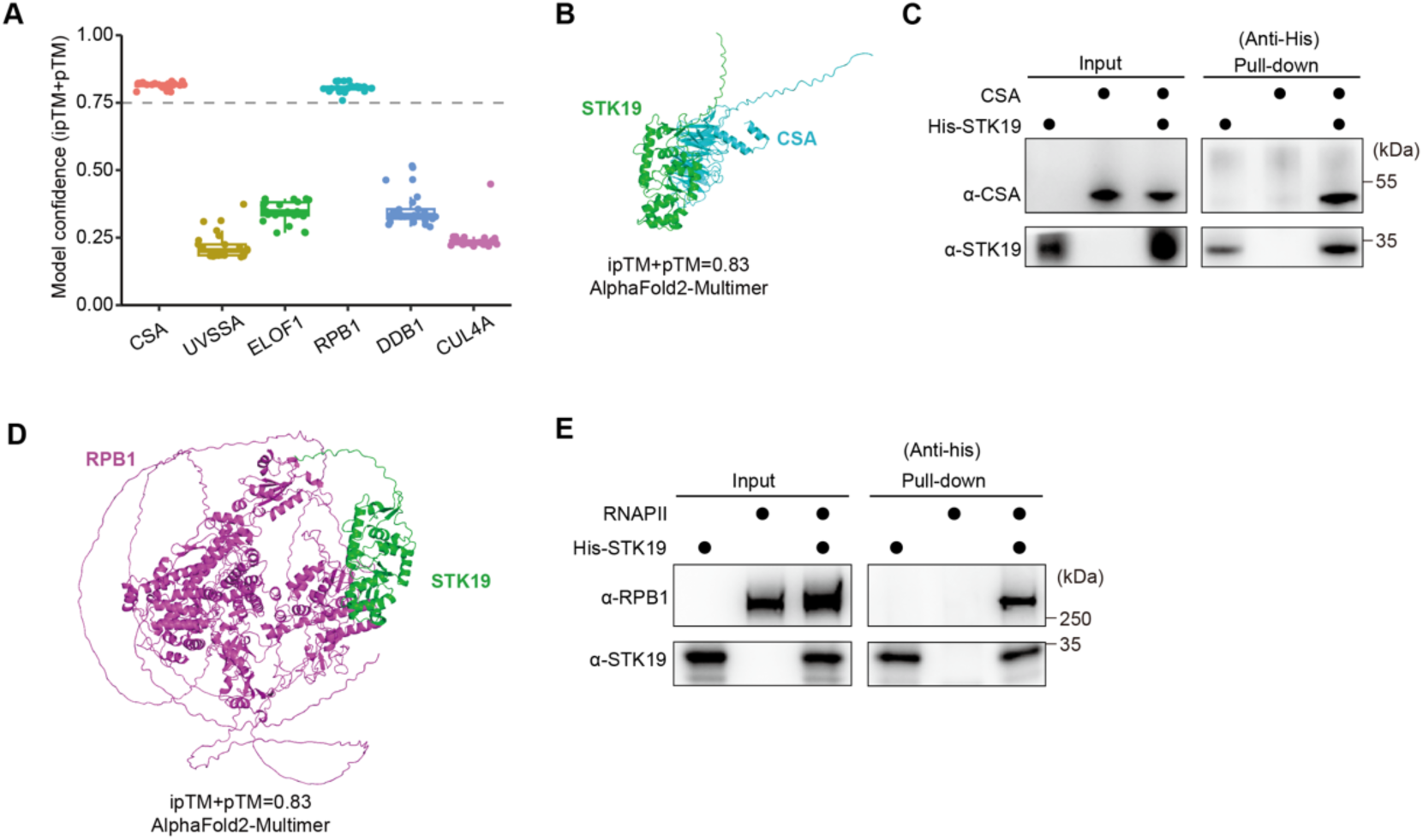
STK19 interacts with CSA and PolII *in vitro*. (A) Prediction of protein interactions between STK19 and known TCR factors by AlphaFold2-Multimer (AF2M). Each prediction generates 5 models and each model contains 5 prediction results. Prediction results with 0.75 or higher model confidence (ipTM+pTM score) are thought to be highly confidential. pTM: predicted Template Modeling, ipTM: interface predicted Template Modeling. (B) The predicted structure showing the interaction between STK19 and CSA by AF2M. The best AF2M-predicted model with highest model confidence is shown. (C) *In vitro* pulldown of CSA by His-STK19 with purified proteins. (D) The predicted structure showing the interaction between STK19 and RPB1 by AF2M. The best AF2M-predicted model with highest model confidence is shown. (E) *In vitro* pulldown of PolII by His-STK19 with purified proteins. See also Supplementary Figure S5.

### STK19 is involved in UV-induced mono-ubiquitination of UVSSA

Since the potential kinase activity of STK19 is not required for TCR, and STK19 has a direct interaction with CSA, we wonder whether STK19 plays a role in the recruitment and/or ubiquitination of other repair factors. Classical TCR factors including CSB, CSA and UVSSA are recruited to chromatin upon UV treatment in TCR-proficient cells, as previously reported(14) and shown by WB (Figure5A-D). Intriguingly, loss of STK19 showed no apparent impact on the recruitment of these factors in both HeLa and XP-C cells (Figure5A-D). However, the upper band in the blot of UVSSA which was reported to be UV-induced UVSSA mono-ubiquitination was impaired in all *STK19*-KO cells (Figure5A-D), similar to *ELOF1*-KO cells (Figure5A)(16,17,19). Accordingly, complement of STK19 could rescue the loss of UVSSA ubiquitination (Figure5B,D). ELOF1 is involved in UV-induced ubiquitination of both UVSSA and RPB1, thus we tested the role of STK19 in RPB1 ubiquitination by DSK2-pulldown assay. DSK2 is a ubiquitin-binding protein that can be used to pull down all ubiquitinated proteins, so the ubiquitination of target protein can be measured by WB of DSK2-pulldown samples. As shown in Figure5E, unlike ELOF1, loss of STK19 did not compromise UV-induced poly-ubiquitination of elongating RPB1. Moreover, recently we found that lesion-stalled PolII can be cleared within 2 h post UV by p97-proteasome pathway in TCR-deficient *UVSSA*-KO cells depending on RPB1 ubiquitination(26). Similar as in *UVSSA*-KO cells, PolII dissociated from damage sites quickly in *STK19*-KO cells although TCR was inhibited (Figure5F), consistent with the unhampered UV-induced RPB1 ubiquitination in this cell line.

**Figure 5.**
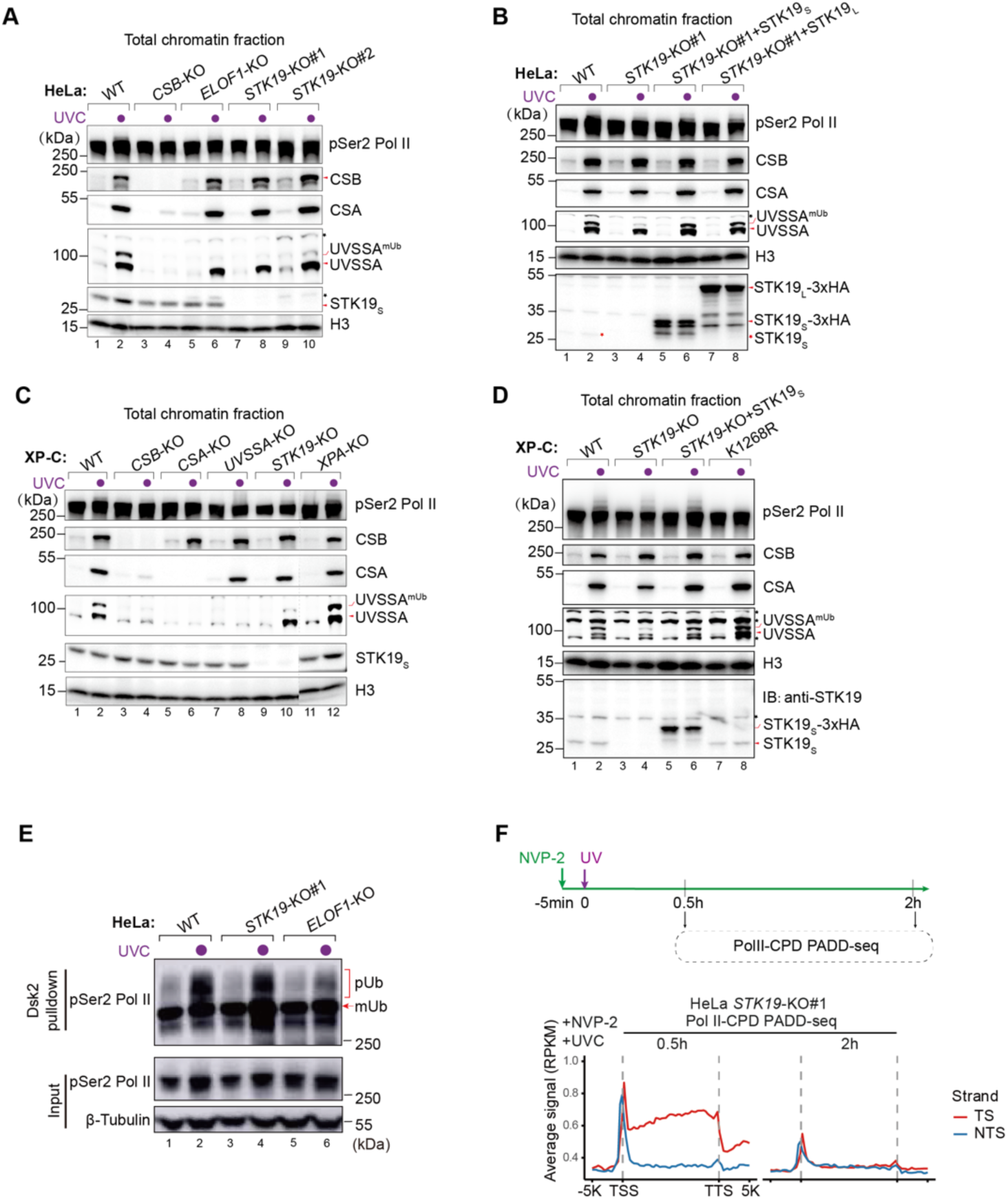
STK19 is involved in UV-induced mono-ubiquitination of UVSSA. (A-D) Western blot analysis showing TCR factors in total chromatin fraction of HeLa-WT, *CSB*-KO, *ELOF1*-KO and *STK19*-KO cells (A), HeLa-WT, *STK19*-KO and STK19-complement cells (B), XP-C Control, *CSB*-KO, *CSA*-KO, *UVSSA*-KO, *STK19*-KO and *XPA*-KO cells (C), XP-C Control, RPB1*-*K1268R, *STK19*-KO and STK19-complement cells (D) at 0.5 h after UVC irradiation for detecting. Non-specific bands are indicated with black asterisk. (E) Detecting UV-induced Pol II polyubiquitylation by Dsk2 pulldown assay in HeLa-WT, *STK19*-KO and *ELOF1*-KO cell lines. (F) Meta-analysis of Pol II-CPD PADD-seq in HeLa *STK19*-KO cells at 0.5 h and 2 h after UVC irradiation. NVP-2 was added to prevent *de novo* PolII release after UV treatment.

### STK19 participates in TFIIH loading and interacts with TFIIH *in vitro*

Our data suggested a nearly complete inhibition of TCR by *STK19* knockout (Figures1,2), thus we wondered whether affecting UVSSA mono-ubiquitination is the sole role of STK19 in TCR. Therefore, we expressed UVSSA-WT and UVSSA-K414R in *UVSSA*-KO XP-C cells and assessed their TCR capacity by Damage-seq (FigureS6A-D). K414 is the main ubiquitination site of UVSSA, thus the UVSSA-K414R mutant protein cannot be ubiquitinated after UV irradiation(19). As shown in FigureS6C,D, UVSSA-WT could efficiently rescue TCR, while UVSSA-K414R also partially recovered TCR, arguing the indispensable role of UVSSA mono-ubiquitination in TCR. Then we checked the impact of STK19 on TCR in UVSSA-K414R mutant cells by siRNA knockdown (Figure6A). Intriguingly, loss of STK19 could further inhibit TCR in UVSSA-K414R mutant cells (Figure6B,C), indicating a UVSSA-K414ub-independent role of STK19 in human TCR.

**Figure 6.**
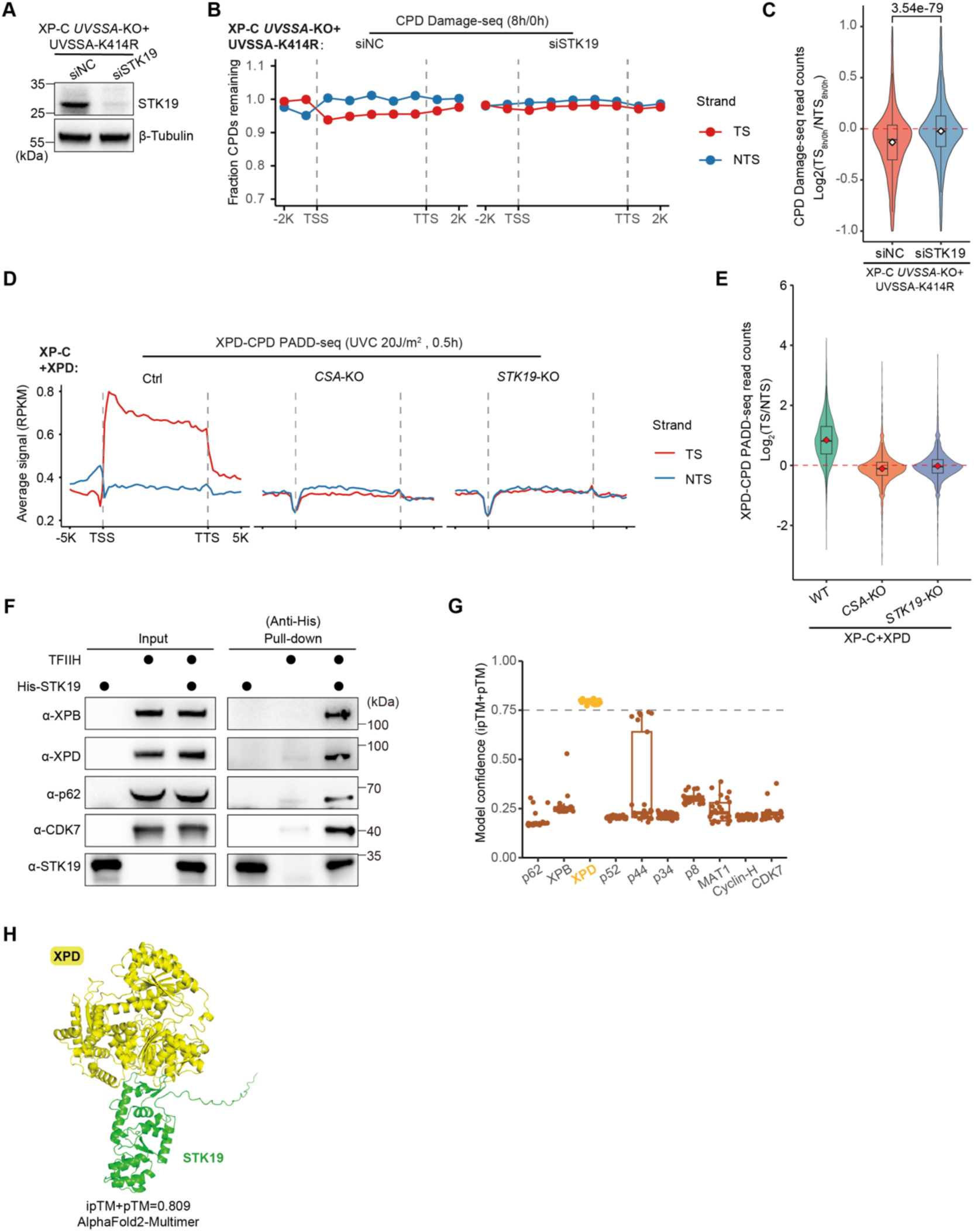
STK19 participates in TFIIH loading and interacts with TFIIH *in vitro*. (A) Western blot analysis showing STK19 knockdown by siRNA in XP-C UVSSA-K414R rescued cells. (B) CPD repair in STK19-depleted or control XP-C UVSSA-K414R cells measured by Damage-seq. The average fraction of CPDs remaining after 8h of repair relative to 0h was plotted along the TSs and NTSs for 9973 nonoverlapping protein-coding genes. (C) Relative quantification of TCR activity in STK19-depleted or control XP-C UVSSA-K414R cells based on the ratio of remaining CPD in TSs to NTSs for 5910 expressed protein-coding genes (TPM > 1). (D) Metaplots of XPD-CPD PADD-seq signal in the indicated XP-C cell lines overexpressing XPD. A total of 4869 expressed nonoverlapping protein-coding genes were used. (E) Relative quantification of XPD-CPD PADD-seq. A total of 6390 expressed protein-coding genes were used. (F) *In vitro* pulldown of TFIIH complex by His-STK19 with purified proteins. (G) Prediction of protein interaction between STK19 and components of TFIIH complex by AlphaFold2-Multimer (AF2M). Each prediction generates 5 models and each model contains 5 prediction results. Prediction results with 0.75 or higher model confidence (ipTM+pTM score) are thought to be highly confidential. (H) The predicted structure showing the interaction between STK19 and XPD by AF2M. The best AF2M-predicted model with highest model confidence is shown. See also Supplementary Figure S6 and S7(A-C).

The recruitment of TFIIH is a key step of TCR, thus we checked the influence of STK19 on TFIIH loading by PADD-seq. HA-tagged XPB or XPD was expressed in XP-C cells (FigureS6E), then the interaction between TFIIH and CPDs was measured by XPB/XPD-CPD PADD-seq. As shown in Figure6D,E and FigureS6F,G, TFIIH bound to CPDs on TSs in parental XP-C cells, while this interaction disappeared in either *CSA*-KO or *STK19*-KO cells, indicating that STK19 is required for TFIIH loading. Therefore, we checked the direct interaction between STK19 and TFIIH, and found that STK19 could pull down the 10-subunit full TFIIH complex *in vitro* (Figure6F, FigureS7A). Among the 10 TFIIH subunits, only XPD was predicted to have a credible interaction with STK19 in all tested models (Figure6G,H, FigureS7B,C), implying the interaction between STK19 and TFIIH is likely through XPD.

## Discussion

Although TCR was first discovered in mammalian cells nearly 40 years ago(56), the molecular mechanism of mammalian TCR still remains unclear. Specifically, whether all essential factors of TCR have been identified is unknown due to the lack of an *in vitro* reconstituted system. STK19 has attracted our attention since its loss can sensitize cells to UV and UV-mimic damage, and compromise the recovery of RNA synthesis after UV treatment(21,22). However, UV damage can trigger global transcription suppression in trans besides directly blocking PolII elongation in cis, both of which are related to cell survival and RNA synthesis(57). As examples, both PAF1C and EXD2 played important roles in cell survival and recovery of RNA synthesis upon UV treatment by stimulating transcription restarting after damage, albeit they were not involved in TCR(58,59). Although the possibility that STK19 is also involved in transcription restarting independent of its role in TCR cannot be excluded, we demonstrated that STK19 is essential for TCR by directly measuring TCR with XR-seq and Damage-seq in HeLa and XP-C cells, respectively. In addition, as demonstrated by PADD-seq, STK19 is recruited to damage sites during TCR, while it is not binding to damage in the absence of CSB or CSA, in contrast to PolII that is tightly restrained on damage under such conditions(37). Therefore, STK19 is more like a classical TCR factor which is recruited to lesion-stalled PolII after damaging rather than a transcription elongation factor that is moving along with PolII even without damage, although it has not been reported to be related to human genetic diseases such as CS and UV^S^S. It is worth noting that STK19 binds to chromatin without damage independent of CSB and CSA (Figure5A,C). This is in line with recent studies reporting that STK19 is a DNA/RNA binding protein(24,60), implying that STK19 has other functions beyond TCR.

According to current knowledge, CSB, CSA (in the form of CRL4^CSA^ ubiquitin E3 ligase) and UVSSA are sequentially recruited to lesion-stalled PolII during TCR, while RPB1 and UVSSA are ubiquitinated by CRL4^CSA^ with the aid of ELOF1(4,5,20). Since the recruitment of STK19 requires CSB and CSA but not ELOF1 and UVSSA, nor ubiquitination of RPB1, it is reasonable to speculate that STK19 is recruited by CSA, which is confirmed by the direct interaction between STK19 and CSA. It is worth noting that STK19 could directly interact with PolII *in vitro*, albeit it cannot be recruited in the absence of CSB or CSA. This interaction might also contribute to the recruitment of STK19. However, subsequent steps of TCR, i.e., the recruitment of UVSSA and ubiquitination of RPB1, do not require STK19, indicating that the recruitment of STK19 is parallel with UVSSA loading and RPB1 ubiquitination. Protein alignment based on reported PolII-ELOF1-CSB-CRL4^CSA^-UVSSA-DNA structure(20) and predicted CSA-STK19 or RPB1-STK19 structure shows no major spatial conflict (Figure7A, FigureS7D). Intriguingly, the key residues of STK19 for DNA binding are close to DNA in the aligned complex (Figure7B), indicating a role of its DNA binding capacity in TCR. Thus, these data suggest that STK19 is a missing piece of the TCR complex. Future study on the real structure of TCR complex containing STK19 is required.

**Figure 7.**
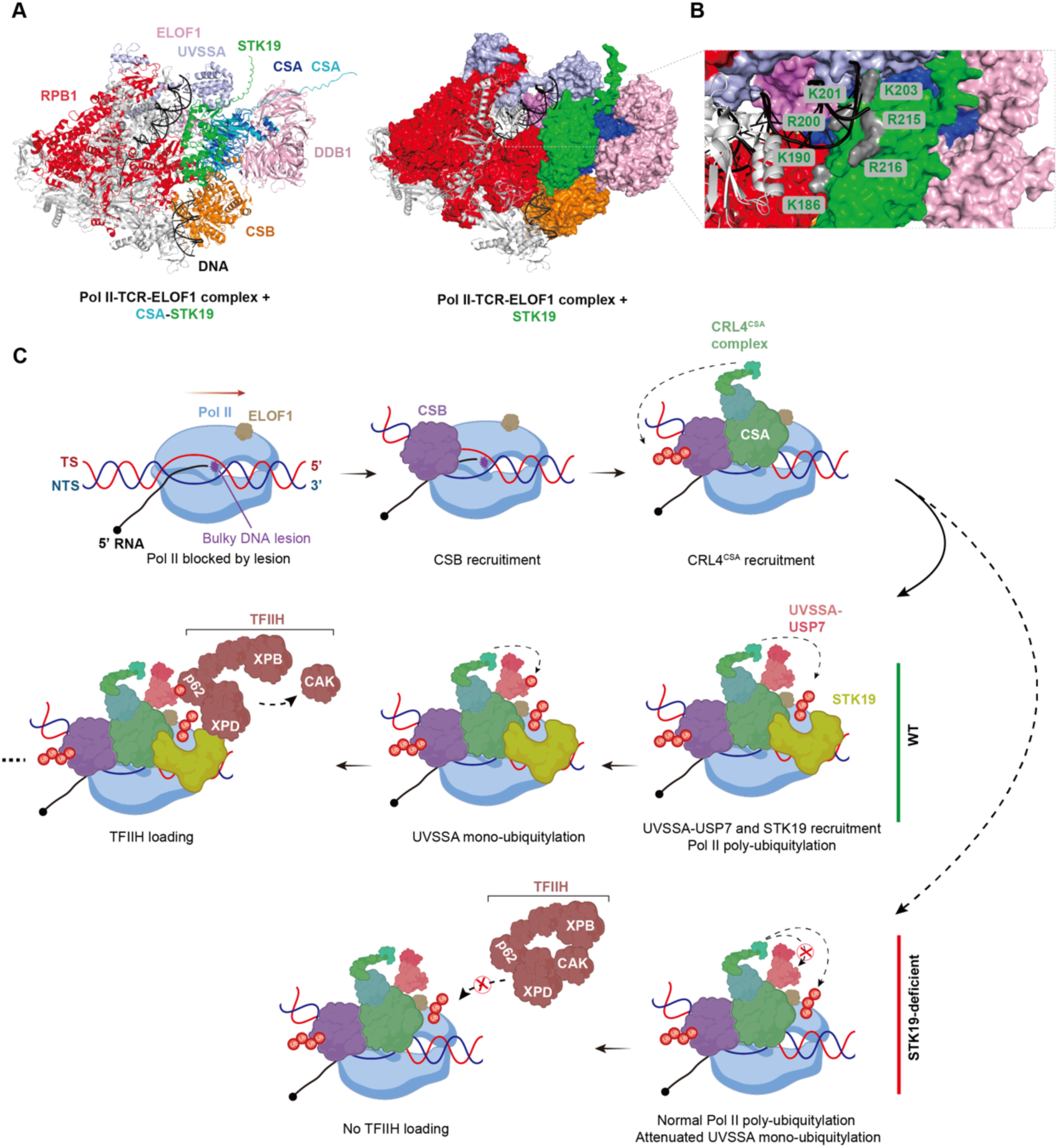
Proposed model of STK19 in TCR. (A) The structural alignment based on experimental TCR complex (PDB: 8B3D) and predicted STK19-CSA complex (best AF2M-predicted model). Left (cartoon), STK19-CSA structure superimposed onto Pol II-TCR-ELOF1 structure; and right (surface), CSA structure from STK19-CSA complex is not displayed. (B) Key residues for STK19 DNA binding capacity (grey)(24,60) are close to DNA in the aligned structure. (C) A proposed model showing the recruitment and essential roles of STK19 in TCR. See also Supplementary Figure S7(D,E).

Our data showed that STK19 can stimulate UVSSA ubiquitination, likely through its binding to CSA. However, why STK19 selectively promotes the ubiquitination of UVSSA but not RPB1 is unclear. Although previous studies suggested that mono-ubiquitination of UVSSA at K414 plays an important role in the recovery of RNA synthesis post UV(19), our results indicated that the ubiquitination-deficient UVSSA-K414R mutant protein could partially rescue TCR in *UVSSA*-KO cells. The fact that depleting STK19 further inhibited residual TCR in UVSSA-K414R mutant cells suggests that STK19 has other role(s) in TCR besides promoting UVSSA ubiquitination. Regarding this question, our TFIIH-CPD PADD-seq results indicate that STK19 is essential for the recruitment of TFIIH. Although the ubiquitination of UVSSA might also affect TFIIH loading, the *in vitro* pull-down assay demonstrated that STK19 can directly bind to the 10-subunit TFIIH complex. It was reported that in GGR, TFIIH is first recruited in the form of 10-subunit complex, then the 3-subunit CAK is discarded and the remaining 7-subunit core complex participates in repair(61). Although the detail of TCR is unclear, the ability of STK19 to bind to 10-subunit TFIIH complex suggests the possibility that STK19 is directly involved in recruiting TFIIH during TCR. Notably, the protein interaction prediction showed that STK19 may interact with the XPD subunit of TFIIH, differing from the p62 subunit which is bound by UVSSA and XPC(62). Therefore, STK19 and UVSSA may collaborate to recruit TFIIH by interacting with different subunits. Based on these results, we proposed a model for TFIIH loading of human TCR involving STK19 (Figure7C). When elongating PolII is blocked by TBLs, CSB and CRL4^CSA^ are sequentially recruited. Then UVSSA and STK19 are recruited and bind to DNA(24,63), while RPB1-K1268 is ubiquitinated by CRL4^CSA^ with the help of ELOF1 and UVSSA(20). STK19, ELOF1 and RPB1-K1268ub can aid the mono-ubiquitination of UVSSA at K414 by CRL4^CSA^(16,19). Finally, TFIIH is recruited through the direct interactions with STK19 (via XPD) and UVSSA (via p62). In a certain sense, STK19 is like an adapter in the TCR complex to connect DNA, PolII, CSA and TFIIH. However, the specific impacts of these interactions in TCR are unknown. Since STK19 is a compact protein and the interfaces of these interactions are interwoven (FigureS7E), it is difficult to abrogate one interaction without affecting others. Further structural and biochemical studies are required to find out key residues involving in these interactions and unveil the function of each interaction in TCR.

Mutations of STK19, especially D89N, was reported to be linked to melanoma in a kinase-dependent manner(23). However, the long isoform of STK19 on which the D89N mutation locates and the potential kinase activity are not required for TCR. Indeed, the CS and UV^S^S patients with TCR deficiency do not show higher risk of skin cancer(64,65). Thus, the role of STK19 in melanoma is unlike to be associated with its TCR function. Since loss of STK19, similar to UVSSA(26), can inhibit TCR without influencing its dissociation from damage sites, we speculate that loss of its TCR function should cause a similar consequence as UVSSA deficiency which results in UV^S^S. However, the incidence of UV^S^S might be substantially underestimated due to its relatively mild symptoms(65). This may explain the fact that no human disease has been reported to be related to the TCR function of STK19 till now. Further study is needed to explore the relationship between the TCR function of STK19 and human health.

## SUPPLEMENTARY DATA

Supplementary Data are available at xxx.

## DATA AVAILABILITY

All raw sequencing data are available at the NCBI Sequence Read Archive (https://www.ncbi.nlm.nih.gov/sra/) with BioProject ID xxxxxx. The codes are publicly available at xxxxx.

## ACKNOWLEDGEMENTS

This work was supported by the Medical Research Data Center of Fudan University. We thank Dr. Feilong Meng (Center for Excellence in Molecular Cell Science, Chinese Academy of Sciences, China) for providing the pMXs-IRES-Puro retroviral expression vector, packaging plasmid PCL10A1, and pGEX-GST-Dsk2 plasmid, and Drs. Yanhui Xu and Xizi Chen (Fudan University, China) for providing the purified human RNA polymerase II and TFIIH complexes.

## FUNDING

This work was supported by National Key R&D Program of China [2022YFA1303000], National Natural Science Foundation of China (NSFC) [32271343], Shanghai Municipal Natural Science Foundation [22ZR1413900], and innovative research team of high-level local university in Shanghai (to J.H.).

## Notes

### Competing Interest Statement

The authors have declared no competing interest.

